# RNase L amplifies Interferon signaling by inducing PKR-mediated antiviral stress granules

**DOI:** 10.1101/2020.02.07.939645

**Authors:** Praveen Manivannan, Mohammad Adnan Siddiqui, Krishnamurthy Malathi

## Abstract

Virus infection leads to activation of the interferon-induced endoribonuclease, RNase L, which results in degradation of viral and cellular RNAs. Both cellular and viral RNA cleavage products of RNase L bind pattern recognition receptors (PRR) like Retinoic acid-inducible I (Rig-I) and or melanoma differentiation-associated protein 5 (MDA5) to further amplify interferon (IFN) production and antiviral response. Although much is known about the mechanics of ligand binding and PRR activation, how the cells coordinate RNA sensing to signaling response and interferon production remains unclear. We show that RNA cleavage products of RNase L activity induce formation of antiviral stress granule (avSG) by regulating activation of double-stranded RNA (dsRNA)-dependent protein kinase R (PKR), and recruit antiviral proteins Rig-I, PKR, OAS and RNase L to avSG. Biochemical analysis of purified avSG showed interaction of key stress granule protein, G3BP1, with only PKR and Rig-I and not with OAS or RNase L. AvSG assembly during RNase L activation is required for IRF3-mediated IFN production and not IFN signaling or proinflammatory cytokine induction. Consequently, cells lacking avSG formation or RNase L signaling produced less IFN and showed higher susceptibility during Sendai virus infection demonstrating the importance of avSG in RNase L-mediated host defense. During viral infection, we propose a role for RNase L-cleaved RNAs in inducing avSG containing antiviral proteins to provide a platform for efficient interaction of RNA ligands with pattern recognition receptors to enhance IFN production to effectively mount antiviral response.

**IMPORTANCE:** Double-stranded RNAs produced during viral infections serve as pathogen associated molecular patterns (PAMPs) and bind pattern recognition receptors to stimulate IFN production. RNase L is an IFN-regulated endoribonuclease that is activated in virus-infected cells and cleaves single-stranded viral and cellular RNAs. The RNase L-cleaved dsRNAs signal to Rig-like helicases to amplify IFN production. This study identifies a novel role of antiviral stress granules induced by RNase L as an antiviral signaling hub to coordinate the RNA ligands with cognate receptors to mount effective host response during viral infections.

## INTRODUCTION

Viral invasion and replication are detected in host cells by Pathogen Recognition Receptors (PRRs) triggering signaling pathways that result in production of type 1 interferon (IFN) (1–3). IFN produced by virus-infected cells acts in autocrine and paracrine ways by binding to cell surface receptors (IFNAR) to induce expression of antiviral IFN-stimulated genes (ISGs) including Rig-I, MDA5, 2’-5’-oligoadenylate synthetase (OAS), RNase L, dsRNA-dependent protein kinase R (PKR), Interferon Induced proteins with Tetratricopeptide repeats (IFIT) to perpetuate antiviral signaling(3, 4). Recognition of viral nucleic acids that serve as Pattern Associated Molecular Patterns (PAMPs) is accomplished by PRRs like the endosomal Toll-like Receptors (TLR3, 7/8, 9), cytosolic Rig-I like receptors (RLRs, Rig-I, MDA5), DExD/H-box helicases (DDX1, DDX21,DHX33, and DHX36), protein kinase R (PKR), 2’,5’-oligoadenylate synthetases (OAS), or cytosolic DNA sensors (DAI, STING, cGAS) by virtue of their compartment-specific distribution in cells and modification on the RNAs or DNA (5–12). RLRs detect cytoplasmic viral RNAs and discriminate self from viral RNAs by recognizing double-stranded structures and 5’-triphosphate that are lacking on self RNAs(13). Rig-I and MDA5 contain an RNA helicase domain for binding RNA and a caspase recruitment domain (CARD) for downstream signaling (14). In case of Rig-I, RNA binding allows K63 ubiquitination of the CARD domain by TRIM25 and ATPase activity induces conformational change and oligomerization (15). In contrast, Rig-I undergoes degradation after conjugation to E3 ubiquitin ligase RNF125 (16). The Rig-I CARD domain interacts with the CARD-like domain of the IFN-β promoter stimulator-1 (IPS-1/MAVS/VISA/Cardif) at the outer mitochondrial membrane via CARD-CARD interaction, which further activates TRAF3 and TBK1(17–20). TBK1 phosphorylates IRF3 that translocate to the nucleus to induce IFN production (21, 22).

Double-stranded RNA-dependent protein kinase R (PKR) is activated by binding dsRNA ligands and participates in integrated stress response during viral infections (23). RNA binding induces dimerization and autophosphorylation resulting in activation and phosphorylation of eIF2α (eukaryotic initiation factor 2 alpha subunit)(24). Phosphorylated eIF2α represses translation and causes aggregation of stalled translation preinitiation complexes containing mRNAs, initiation factors, small ribosomal subunits, RNA-binding proteins together with the Ras-GAP SH3 domain binding protein (G3BP) and T-cell restricted intracellular antigen 1(TIA 1) into stress granules (SG)(25).

SGs are nonmembranous RNA-protein complexes that are formed in the cytoplasm in response to diverse stress signals including viral infections (26). Complex RNA-protein interactions in the SG establish a liquid-liquid phase separation from the rest of the cytoplasm, facilitating recruitment of multiple proteins into a dynamic compartment (27). Depending on the nature of the stress signal protein kinases such as PKR (protein kinase R), GCN2 (general control nonderepressible 2), HRI (heme-regulated inhibitor) or PERK (PKR-like ER kinase) phosphorylate translation initiation factor, eIF2α, to inhibit translation which in turn promotes SG formation (28–30). SG composition and proteins recruited vary depending on the type of stimulus and cell type but G3BP1 is required for nucleation in all contexts. SG were considered general triage sites for mRNA turnover, however, recent studies show selective exclusion from SGs of some transcripts needed to overcome stress (31–33). Compared to these canonical SGs formed under stress conditions, antiviral SGs form during viral infections and have been proposed to play a role in antiviral signaling (34) by recruiting antiviral proteins including PKR, Rig-I, MDA5, OAS, RNase L, Trim5, ADAR1, ZAP, cGAS and RNA helicases like DHX36, DDX3 and DDX6 (35–38). Assembly of avSG was required for signaling to produce IFN during NDV, IAV and SINV infections. During IAV infection, SG are induced and both IAV viral RNA and Rig-I are sequestered in SG, thereby providing a platform for sensing of viral RNA by Rig-I (35, 39). In addition, antiviral proteins like PKR, OAS, RNase L, LGP2 and MDA5 were also shown to coalesce in avSGs (35, 36, 40, 41). Several viruses counteract SG formation by targeting SG proteins through G3BP1 cleavage or by inhibiting upstream eIF2α pathway to support viral replication suggesting an important role of SG in viral pathogenesis (42, 43).

In most vertebrates, viruses induce an RNA degradation process that is regulated through the action of the ubiquitous cellular endoribonuclease, RNase L. Type I IFN produced during viral infections transcriptionally induces OAS proteins that are activated by binding dsRNA to produce a unique 2’, 5’-oligoadenylate, 2-5A ((p*x*5′A(2′p5′A)*n*; *x* = 1–3; *n* ≥ 2), produced from cellular ATP. The only established function of 2-5A is activation of RNase L. RNase L is expressed as an inactive monomer and binding 2-5A promotes dimerization and conversion to an active enzyme that targets single-stranded viral and cellular RNA after UU/UA nucleotide sequences resulting in dsRNA cleavage products with 5’-hydroxyl and 2’,3’-cyclic phosphate ends(10, 44, 45). While activity of RNase L on viral genome or mRNA directly eliminates viruses, the dsRNA cleavage products signal through Rig-I/MDA5/ MAVS (IPS-1) and IRF3 to induce IFN production (46). RNase L-cleaved RNAs also induce NLRP3 inflammasome and promote switch from RNase L-induced autophagy to apoptosis by promoting cleavage of autophagy protein, Beclin-1 (47, 48). The role of RNase L in generating dsRNA with IFN-inducing abilities and the multiple overlapping signaling pathways activated by avSG during viral infections prompted us to explore the role of RNase L-cleaved RNAs in inducing avSG formation as a platform for antiviral signaling. Our results show that direct activation of RNase L with 2-5A or treatment with RNase L-cleaved RNAs induced avSG formation by activating PKR and phosphorylation of eIF2α. Characterization of purified avSG showed the interaction of G3BP1 in avSG with only PKR and Rig-I, but not OAS or RNase L. AvSG assembly was required for IRF3-mediated IFN production, but not IFN signaling or proinflammatory cytokine induction and affected viral pathogenesis. These studies demonstrate avSG assembly induced by RNase L as an antiviral signaling hub to coordinate RNA ligands with PRRs to mount effective antiviral response.

## RESULTS

### RNase L activation induces formation of antiviral stress granules containing antiviral proteins

Viral infection or dsRNA causes aggregation of Rig-I, PKR, OAS and RNase L in antiviral stress granules (avSG) for production of type I IFN by providing a platform for integrating RNA ligands with antiviral proteins. The role of dsRNA by-products of RNase L enzyme activity in regulating type I IFN production prompted us to explore avSG formation during RNase L activation. Transfection of HT1080 fibrosarcoma cells with 2-5A, a highly specific ligand and activator of RNase L, resulted in a characteristic rRNA cleavage pattern (Fig. 1A) and localization of key stress granule protein, G3BP1, in distinct stress granules compared to diffuse distribution in mock treated cells (Fig. 1B). To determine if these 2-5A-induced stress granules are antiviral stress granules, we performed immunofluorescence assays for RNA-binding antiviral proteins Rig-I, PKR, OAS and RNase L with G3BP1. We observed significant co-localization of these antiviral proteins on RNase L activation in avSGs (Fig. 1B). Following 2-5A transfection, compared to mock treated cells, 32% cells formed stress granules (Fig. 1C). To demonstrate that avSGs were formed in response to RNase L activation, we generated CRISPR-mediated knockout of RNase L in HT1080 cells (49) and observed no avSG formation with 2-5A transfection (Fig.1D). AvSG formation was restored in these cells only by expression of Flag-WT RNase L and not Flag-RNase L R667A mutant which lacked enzyme activity (Fig. 1D) (Ref). These results suggest that direct activation of RNase L by 2-5A induces formation of avSG and Rig-I, PKR, OAS and RNase L are recruited to these avSG.

**Figure 1.**
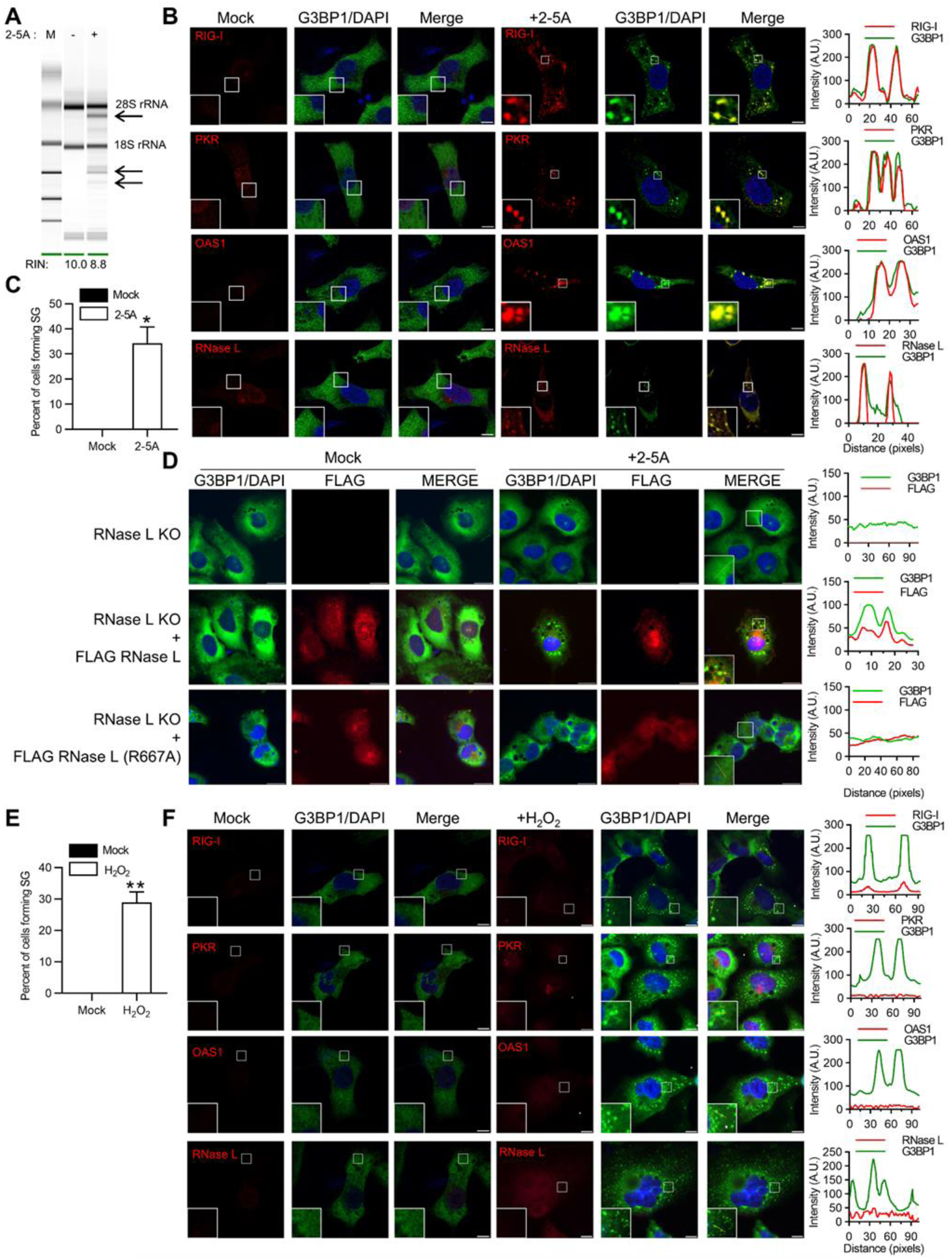
Activation of RNase L induces antiviral stress granules formation. HT1080 cells were transfected with 2–5A (10 µM) for 8h and (A) RNase L-mediated cleavage of rRNA (arrows) was analyzed on RNA chips using the Agilent Bioanalyzer 2100, RNA Integrity Number (RIN) is shown, (B) cells were fixed and stained with G3BP1 and indicated antiviral proteins, the magnified images correspond to the boxed regions, (right) intensity profiles of G3BP1 and antiviral proteins along the plotted lines as analyzed by Image J line scan analysis and (C) the percentage of cells forming stress granules were quantitated. (D) RNase L KO cells were either mock transfected or transfected with FLAG-WT-RNase L or FLAG-R667A-RNase L and immunostained for G3BP1 and FLAG, (right) intensity profiles of G3BP1 and FLAG along the plotted lines as analyzed by Image J line scan analysis. HT1080 cells were treated with H_2_O_2_ (1 mM) for 3h and (E) the percentage of cells forming stress granules were quantitated, (F) cells were immunostained with G3BP1 and indicated antiviral proteins (right) intensity profiles of G3BP1 and antiviral proteins along the plotted lines as analyzed by Image J line scan analysis. All experiments included at least 100 cells from three replicates. Scale bars correspond to 10µm. Data are representative of at least three independent experiments. *p<0.01, **p<0.001

Antiviral stress granules are distinct from canonical stress granules and characterized by the presence of antiviral RNA-binding proteins, RNA helicases and RNA ligands that form during viral infection (50). To demonstrate the distinct nature of avSG, we treated HT1080 cells with H_2_O_2_ to induce oxidative stress that promotes formation of stress granules. While SGs formed in 30% of H_2_O_2_-treated cells as shown by G3BP1 puncta, there was no co-localization of antiviral proteins Rig-I, PKR, OAS or RNase L in these granules (Fig. 1E, F). These results support observations made by others and demonstrate that avSGs are unique and distinct from canonical SGs that form in response to diverse stress stimuli including oxidative stress (35, 36).

### RNase L-cleaved small RNAs activate PKR to induce avSG

The RNA cleavage products of RNase L are predominantly small dsRNA that signal via Rig-I and/or MDA5 and MAVS (IPS-1) to amplify IFN signaling (46). The stress-induced eIF2α kinases like PKR, PERK, GCN2 and HRI phosphorylate eIF2α resulting in formation of SG. PKR is activated by binding dsRNA so we tested the hypothesis that RNase L-cleaved small RNAs activate PKR and promote the formation of avSG by phosphorylation of eIF2α. RNase L-cleaved small RNAs or control small RNAs were purified as previously described (46) and phosphorylation of PKR was monitored following transfection at indicated times in HT1080 cells. RNase L-cleaved small RNAs induced autophosphorylation of PKR 4h post transfection that increased over time compared to control small RNAs (Fig. 2A). PKR phosphorylation correlated with eIF2α phosphorylation only in cells treated with RNase L-cleaved small RNAs but not control small RNAs (Fig. 2B). No phosphorylation of eIF2α by RNase L-cleaved RNAs was observed in cells lacking PKR generated by CRISPR-Cas9 technology (Fig. 2B, C). Taken together, these results indicate that RNase L-cleaved small RNAs activate PKR to phosphorylate eIF2α. To determine if PKR-induced phosphorylation of eIF2α translates into avSG formation, we analyzed avSG formation in PKR KO cells by immunofluorescence analysis and compared with G3BP1 KO, RNase L KO, Rig-I KO and PKR/Rig-I double KO (DKO) cells and calculated the frequency of avSG (Fig. 2D, E). As expected, about 35% of 2-5A transfected WT and Rig-I KO cells formed avSG, and cells lacking RNase L, PKR or PKR/Rig-I DKO did not form avSG. G3BP1 is necessary to form avSG in response to RNase L activation as cells lacking G3BP1 did not induce avSG. When the RNase L-cleaved RNAs were introduced in cells, in addition to WT (38%) and Rig-I KO cells (39%), cells lacking RNase L also formed avSG (34%) suggesting a role for the RNase L cleaved RNAs products in promoting avSG formation. Control small RNAs did not induce avSG in any of the cells. Previous studies show the presence of 5’OH and 2’,3-cyclic phosphoryl on RNase L-cleaved products contribute to IFN production as removal of the 2’,3-cyclic phosphates by treatment with calf intestinal phosphatase reduced IFN production (46). Removal of terminal 2’,3-cyclic phosphoryl on RNase L-cleaved RNAs significantly reduced avSG formation (Fig. 2D, E). In contrast, formation of canonical SGs in response to oxidative stress by H_2_O_2_ is not impacted in cells lacking PKR, RNase L, Rig-I or both PKR/Rig-I (Fig. 2F). Finally, 2-5A transfected cells stained with monoclonal antibody against dsRNA showed co-localization with G3BP1 in avSGs by immunofluorescence assays (Fig. 2G). These findings demonstrate that RNase L enzyme activity produces small dsRNAs that activate PKR to phosphorylate eIF2α and induce formation of avSGs.

**Figure 2.**
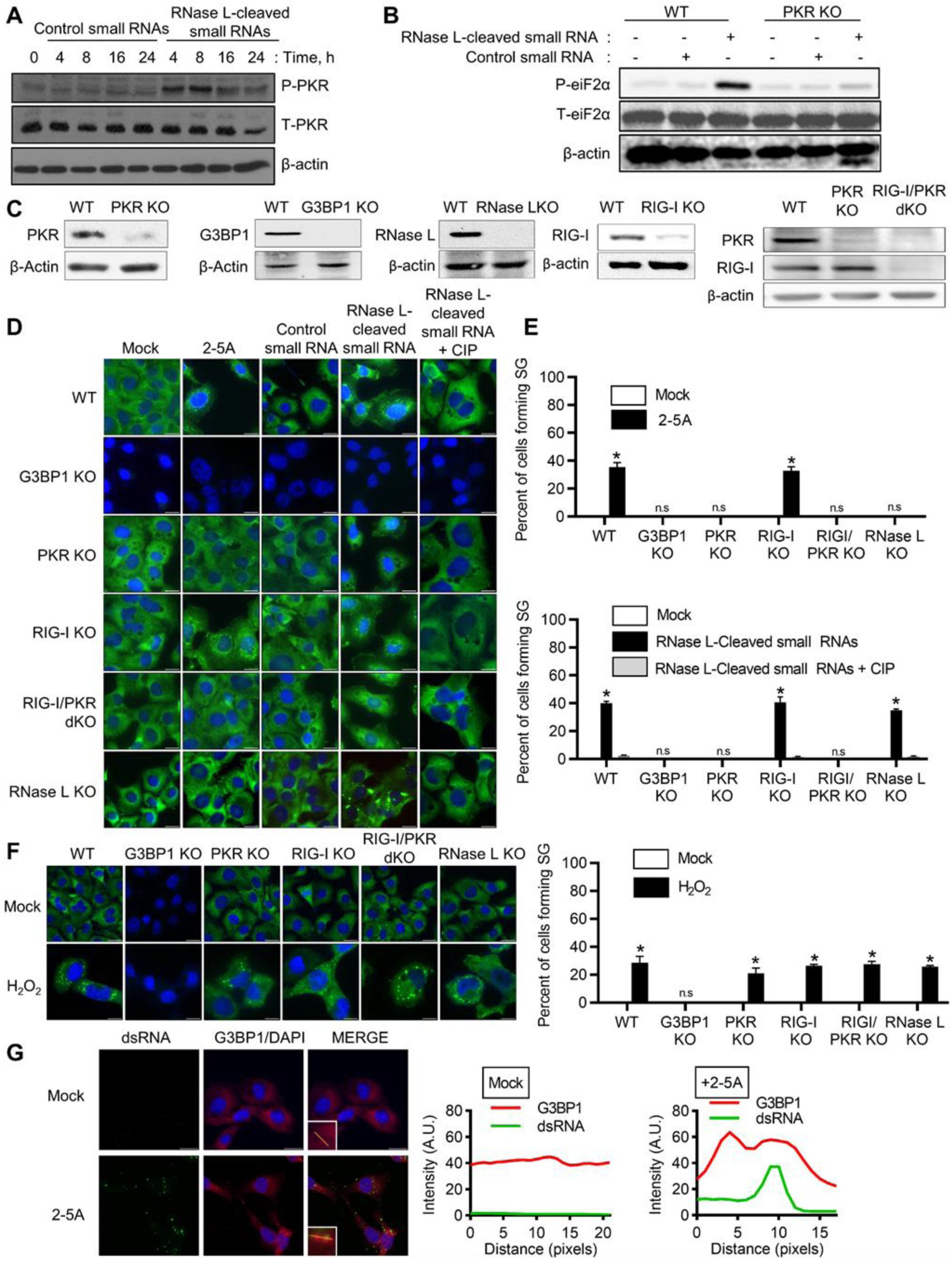
Involvement of PKR in RNase L-mediated avSG formation. (A) HT1080 cells were treated with RNase L-cleaved small RNAs or control small RNAs (2ug/ml) and phosphorylation of PKR was detected in immunoblots, (B) WT and PKR KO cells were treated with control small RNAs or RNase L-cleaved small RNAs (2µg/ml) for 8h and phosphorylation of eIF2α levels were determined in immunoblots, (C) CRISPR/Cas9 knock-out of G3BP1, RIG-I, PKR, RIG-I/PKR or RNase L was verified in cell lysates by immunoblotting using specific antibodies, (D) Indicated cells were treated with 10µM of 2-5A, 2µg/ml of control small RNAs, 2µg/ml RNase L-cleaved small RNAs or 2µg/ml of CIP treated RNase L-cleaved small RNAs for 8h, (E) the percentage of cells forming stress granules were quantitated. (F) Indicated cells were treated with 1mM H_2_O_2_ for 3 hours and stress granule formation analyzed by staining for G3BP1, and the percentage of cells forming stress granules were quantitated. (G) HT1080 cells were treated with 2-5A (10 µM) for 8h and cells were fixed and immunostained with G3BP1 and dsRNA, (right) intensity profiles of G3BP1 and dsRNA along the plotted lines as analyzed by Image J line scan analysis. All experiments included at least 100 cells from three replicates. Data are representative of three independent experiments. Scale bars are 20µm. *p<0.01, n.s: not significant, WT: Wild-Type.

### Co-localization of PKR, Rig-I, OAS and RNase L in avSG on RNase L activation

Studies have reported that avSG provide a platform to coordinate viral sensing and IFN production by recruiting antiviral proteins and RNA ligands. We characterized avSGs formed during RNase L activation by adapting a method recently used to purify SG core from GFP-G3BP1 expressing cells with some changes(51). Expression of GFP-G3BP1 induced SG independent of stimuli, so we used antibodies towards endogenous G3BP1 to immunoprecipitate and test interaction of antiviral proteins from purified avSG core following 2-5A treatment in WT, G3BP1 KO and RNase L KO cells (Fig. 3A, B). As PKR, Rig-I, OAS and RNase L co-localized with G3BP1 on 2-5A treatment, we examined if they purified with avSG core and tested physical interaction in the avSG core by co-immunoprecipitation. Cells were mock transfected or transfected with 2-5A and avSG core were purified by multiple rounds of centrifugation as described in methods. The cell pellet and core were analyzed for expression of PKR, Rig-I, OAS and RNase L and their interaction with G3BP1 was monitored in immune complexes by immunoblotting. Interaction of PKR and Rig-I with G3BP1 in the avSG core was seen only after 2-5A transfection, however, OAS and RNase L were present in the avSG core but did not interact with G3BP1. MAVS (IPS-1), a mitochondrial adaptor protein required for IFN production and RNase L-mediated IFN induction was not present in avSG core (Fig. 3B). As expected, cells lacking G3BP1 or RNase L did not assemble avSG core in response to 2-5A. These results are consistent with avSG co-localization and interaction of G3BP1 with PKR and Rig-I during IAVΔNS1 and NDV infection (35, 36). To further characterize the avSG formation by RNase L-cleaved small RNAs, avSG core was purified from transfected cells and compared to control small RNAs (Fig. 3C). Consistent with avSG formation, RNase L-cleaved small RNAs promoted interaction of G3BP1 with PKR and Rig-I as observed with 2-5A, however, both OAS and RNase L while present in the core do not interact with G3BP1. As expected, when RNase L-cleaved RNAs were introduced into RNase L KO cells, they induced formation of avSG and both PKR and Rig-I interacted with G3BP1 in avSG core and cells lacking G3BP1 did not form avSG (Fig. 3C). We compared these avSG to the canonical SG formed in response to oxidative stress by treating cells with H_2_O_2_ and analyzed the SG core for the presence of antiviral proteins. Consistent with our data in Fig. 2, WT cells formed SG core and none of the antiviral proteins were interacting or present in the SG and cells lacking G3BP1 did not form SG core (Fig. 3D). These results allowed us to analyze the biochemical features of avSG and reveal that unlike antiviral proteins OAS and RNase L that are components of avSG and do not interact with the key proteins like G3BP1, PKR and Rig-I interact with G3BP1 and presumably form the scaffold and core of SG. These results also raise the possibility of existence of different types of complexes in the SG cores that coalesce to form a mature stress granule.

**Figure 3.**
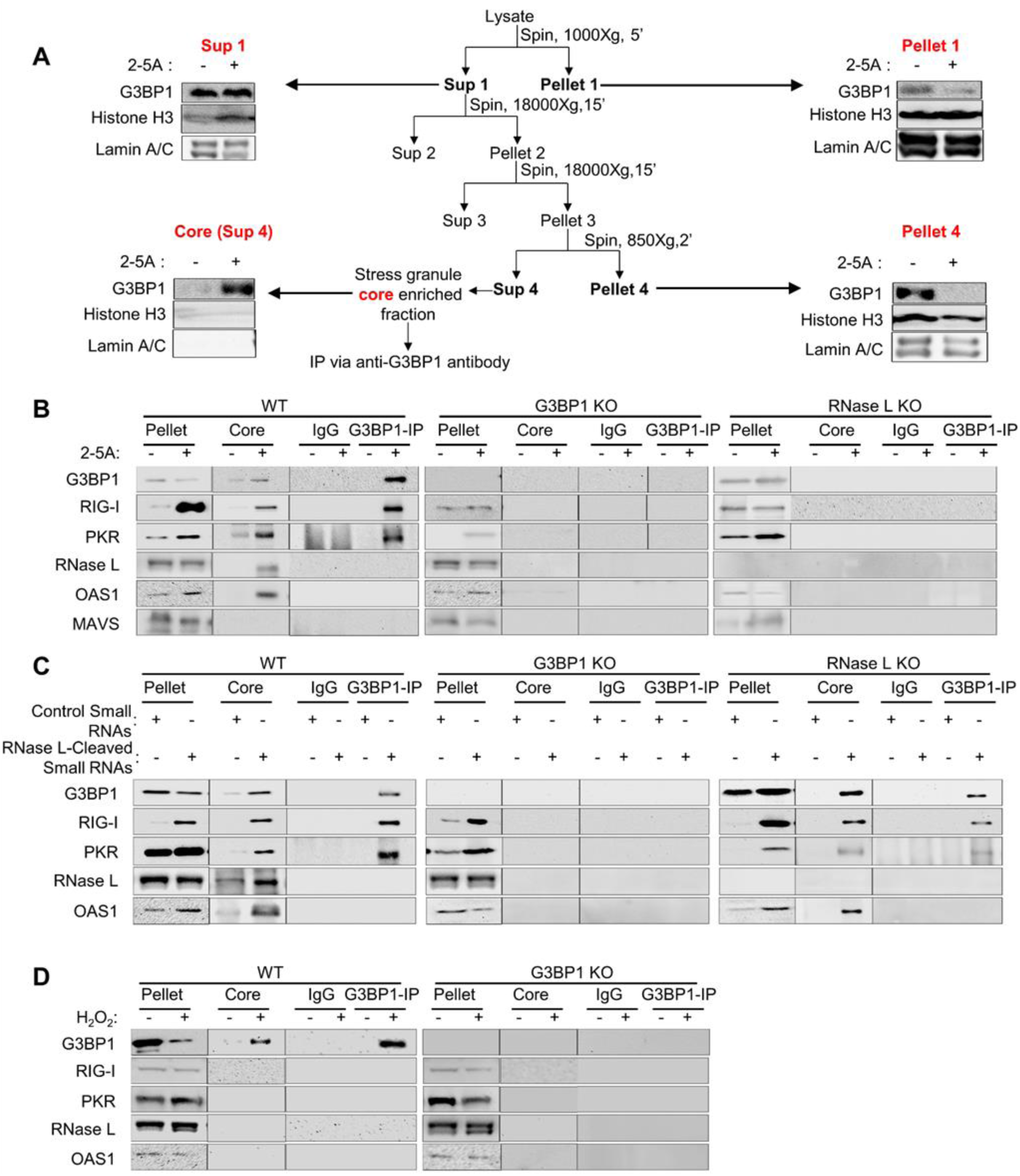
G3BP1 interacts with PKR and Rig-I in avSG, but not OAS and RNase L. (A) Schematic of avSG purification and analysis of fractions in immunoblots using indicated antibodies. HT1080 WT, G3BP1 KO or RNase L KO cells were treated with (B) mock or 10µM 2-5A, (C) RNase L-cleaved small RNAs or control small RNAs (2µg/ml), or (D) 1mM H_2_O_2_ or mock treated and avSG was isolated as described in methods. The avSG core proteins were immunoprecipitated with G3BP1 antibody and immune complex analyzed for presence of PKR, Rig-I, OAS, RNase L and MAVS (IPS-1) by immunoblot analysis. Pellet and SG core fractions were probed for expression of G3BP1 and PKR, Rig-I, OAS, RNase L and MAVS (IPS-1). Nonspecific lanes were cropped to generate the image and the boundaries are indicated. Data are representative of results from two experiments. WT: Wild-Type.

### avSG assembly by RNase L is required for IRF3-mediated interferon induction but not for interferon signaling

Activation of RNase L by 2-5A produces dsRNA intermediates that signal through Rig-I and or MDA5 via mitochondrial adaptor, MAVS (IPS-1) by activating IRF3 that translocates to the nucleus to enhance IFN-β production (46). Various studies have shown that G3BP1 binds to Rig-I to regulate IFN-β production in response to viral RNA and synthetic dsRNA, polyI:C (39, 52, 53). To examine whether G3BP1 participates in RNase L-mediated IFN-β production, we monitored IFN-β promoter activation in WT and G3BP1 KO cells by directly activating RNase L with 2-5A or introducing RNase L-cleaved small RNAs and compared to control small RNAs. In cells lacking G3BP1, IFN-β promoter activation was significantly reduced in response to both 2-5A and RNase L-cleaved small RNAs (Fig. 4A). Consequently, activation of promoters of interferon-stimulated genes (ISGs) like ISG15 and ISG56/IFIT1 that are induced transcriptionally by IFN was also reduced. Consistent with the above observations, mRNA levels of IFN-β, ISG15 and ISG56/IFIT1 were reduced in cells lacking G3BP1 in response to RNase L activation (Fig. 4B). Overexpression of MAVS (IPS-1) activates signaling pathways downstream of Rig-like receptors resulting in phosphorylation and nuclear translocation of IRF3 to promote IFN-β production (18). In the absence of G3BP1, no difference in IFN-β promoter activation or IFN-β mRNA levels was observed in MAVS overexpressing cells suggesting avSG functions upstream of MAVS (IPS-1) (Fig. 4C). This is consistent with the absence of MAVS (IPS-1) in avSG core with RNase L activation (Fig. 3B). Furthermore, overexpression of MAVS (IPS-1) did not induce SG formation (Fig.4C). To further investigate the requirement of avSG in IRF3 activation, we monitored nuclear translocation of GFP-IRF3 in response to 2-5A for indicated times in WT and G3BP1 KO cells by confocal microscopy. Loss of G3BP1 diminished GFP-IRF3 nuclear translocation 3-fold compared to WT cells (29% vs 60%) (Fig. 4D). We used a luciferase-based IRF3 transactivation assay to measure phosphorylation-dependent IRF3 activity. The assay uses Gal4 DNA-binding domain and IRF3 transactivation domain driving luciferase expression under Gal4 promoter when IRF3 is phosphorylated (21). In G3BP1 KO cells, 2-5A induction of IRF3 transactivation was 44% that of WT cells expressing G3BP1 (Fig. 4E). As expected, overexpression of MAVS (IPS-1) resulted in similar levels of IRF3 transactivation independent of G3BP1 expression (Fig. 4F). The effect of G3BP1 on RNase L-mediated IFN-β production was apparent from reduced phosphorylation of PKR, IRF3 and STAT1 following 2-5A treatment in lysates of cells lacking G3BP1 or RNase L compared to strong activation in control WT cells (Fig. 4G). While activation of dsRNA signaling pathway specifically promotes avSG assembly to induce IFN, H_2_O_2_ treatment forms SGs and does not activate PKR, IRF3 or produce IFN (Fig. 4H). Together, our data suggest that G3BP1 is essential for RNase L-mediated IFN induction by promoting avSG assembly containing antiviral proteins and activating IRF3.

**Figure 4.**
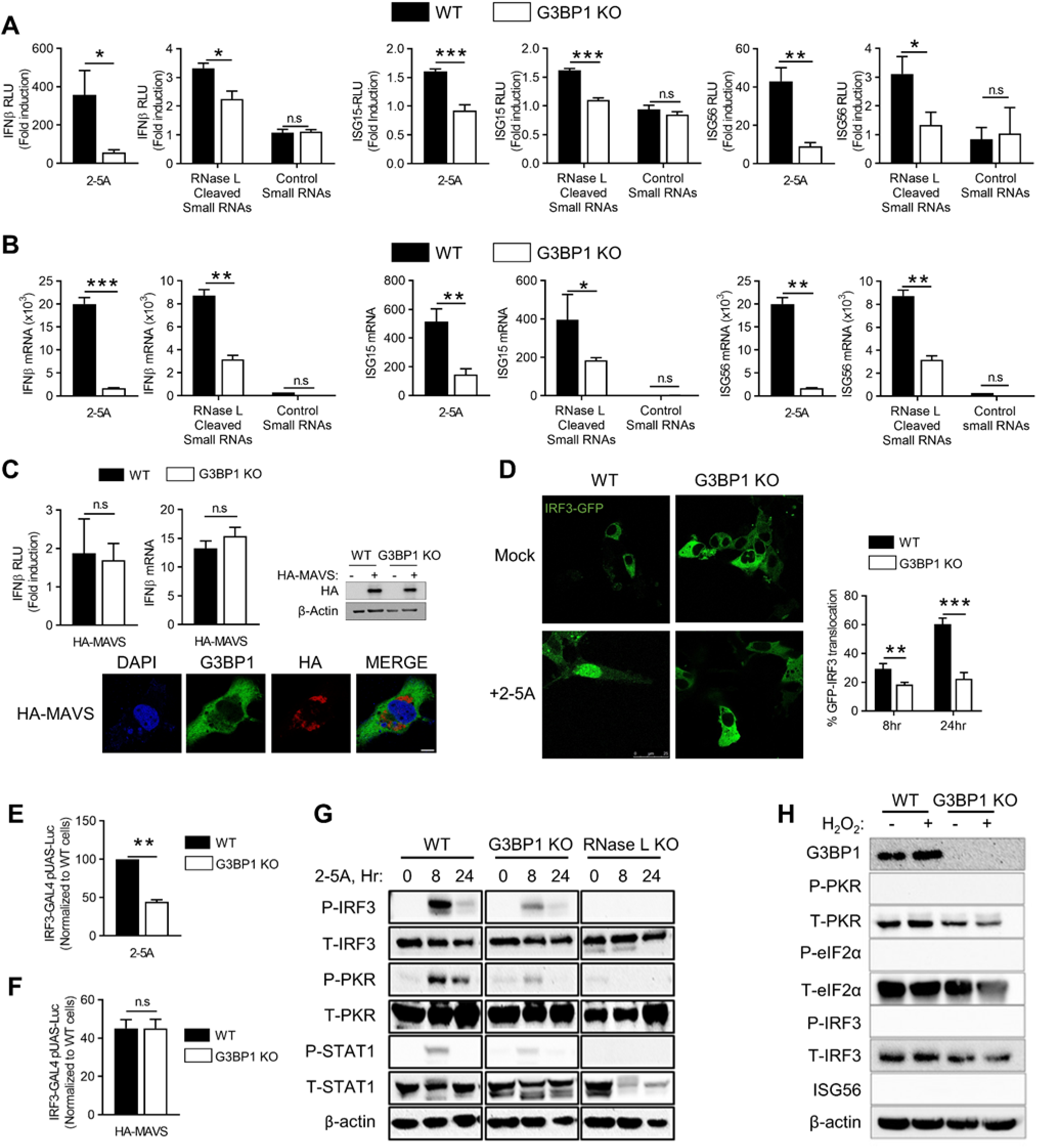
Antiviral SGs are required for IRF3-mediated IFN induction. (A) HT1080 WT and G3BP1 KO cells (1×10^5^) were transfected with IFN-β-luc, ISG15-luc or ISG56-luc reporter constructs along with β-galactosidase plasmids. After 24h, cells were treated with 10µM of 2-5A, 2µg/ml of RNase L-cleaved small RNAs or control small RNAs and 8h later luciferase activity was measured and normalized to β-galactosidase levels. (B) HT1080 WT and G3BP1 KO cells were treated with 10µM of 2-5A, 2µg/ml of RNase L-cleaved small RNAs or control small RNAs and 8h later mRNA levels of IFN-β, ISG15 and ISG56 was measured by qRT-PCR and normalized to GAPDH mRNA levels. (C) WT and G3BP1 KO cells were transfected with empty vector or HA-MAVS(IPS-1), IFN-β-luc and β-galactosidase plasmids and after 24h, promoter activity was normalized to β-galactosidase levels. Effect of HA-MAVS on IFN-β mRNA levels were determined by qRT-PCR and HA-MAVS expression was confirmed in immunoblots. HA-MAVS expressing cells were stained with G3BP1 to determine SG formation. (D) WT and G3BP1 KO cells were transfected with IRF3-GFP and 24h later cells were treated with 10µM of 2-5A or mock treated and imaged after 8h. The percentage of cells with nuclear GFP-IRF3 were calculated in random fields from a minimum of 100 cells and representative images are shown. WT and G3BP1 KO cells were transfected with IRF3-GAL4 and UAS-luciferase plasmids and treated with (E) 10µM of 2-5A for 8h or (F) HA-MAVS. Cells were lysed and luciferase activity was measured. (G) WT, G3BP1 KO or RNase L KO cells were transfected with 10µM of 2-5A for indicated times and p-IRF3, p-PKR and p-STAT1 levels were determined in immunoblot and compared to unphosphorylated levels, β-actin was used to normalize loading. (H) WT and G3BP1 KO cells were transfected with 1mM H_2_O_2_ for 3h and levels of p-PKR, p-eIF2α, p-IRF3 were compared to unphosphorylated levels and induction of ISG56 were determined in immunoblots. Data represent mean ± S.E. for three independent experiments. *p<0.01, **p<0.001, ***p<0.0001, n.s: not significant, WT: Wild-Type.

IFN secreted by virus infected cells binds to type I IFN receptor on cell surface and activates JAK-STAT signaling pathway leading to transcriptional induction of several interferon stimulated genes (ISGs) with roles in viral clearance mechanisms. Our results show the requirement of avSG in producing IFN in response to RNase L activation, however, the role in IFN signaling is not clear. To examine this, we treated cells with type I IFN and monitored transcriptional induction of ISG15 and ISG56 using real-time PCR and promoter-driven luciferase reporter assays. In the absence of G3BP1, no significant differences in mRNA levels of both ISG15 and ISG56 or promoter-driven luciferase activity were observed on IFN treatment (Fig. 5A, B). Exposure of WT, G3BP1 KO or RNase L KO cells to type I IFN resulted in similar levels of phosphorylation of STAT1 accompanied by comparable levels of induction of ISGs like OAS2, OAS3, ISG56 in cell lysates on immunoblots analysis (Fig. 5C). Phosphorylated STAT1 translocates to the nucleus to induce transcription of genes regulated by IFN-stimulated response elements (ISRE) (3). No significant difference in phospho-STAT1 accumulation in the nucleus was observed with IFN treatment in cells lacking G3BP1 or RNase L compared to control WT cells (Fig. 5C, D). Taken together, these results indicate that avSG assembly that requires G3BP1 protein, is required for IFN production in response to RNase L activation. However, following IFN production, G3BP1 is dispensable for activation of JAK-STAT signaling pathway to transcriptionally induce ISGs.

**Figure 5.**
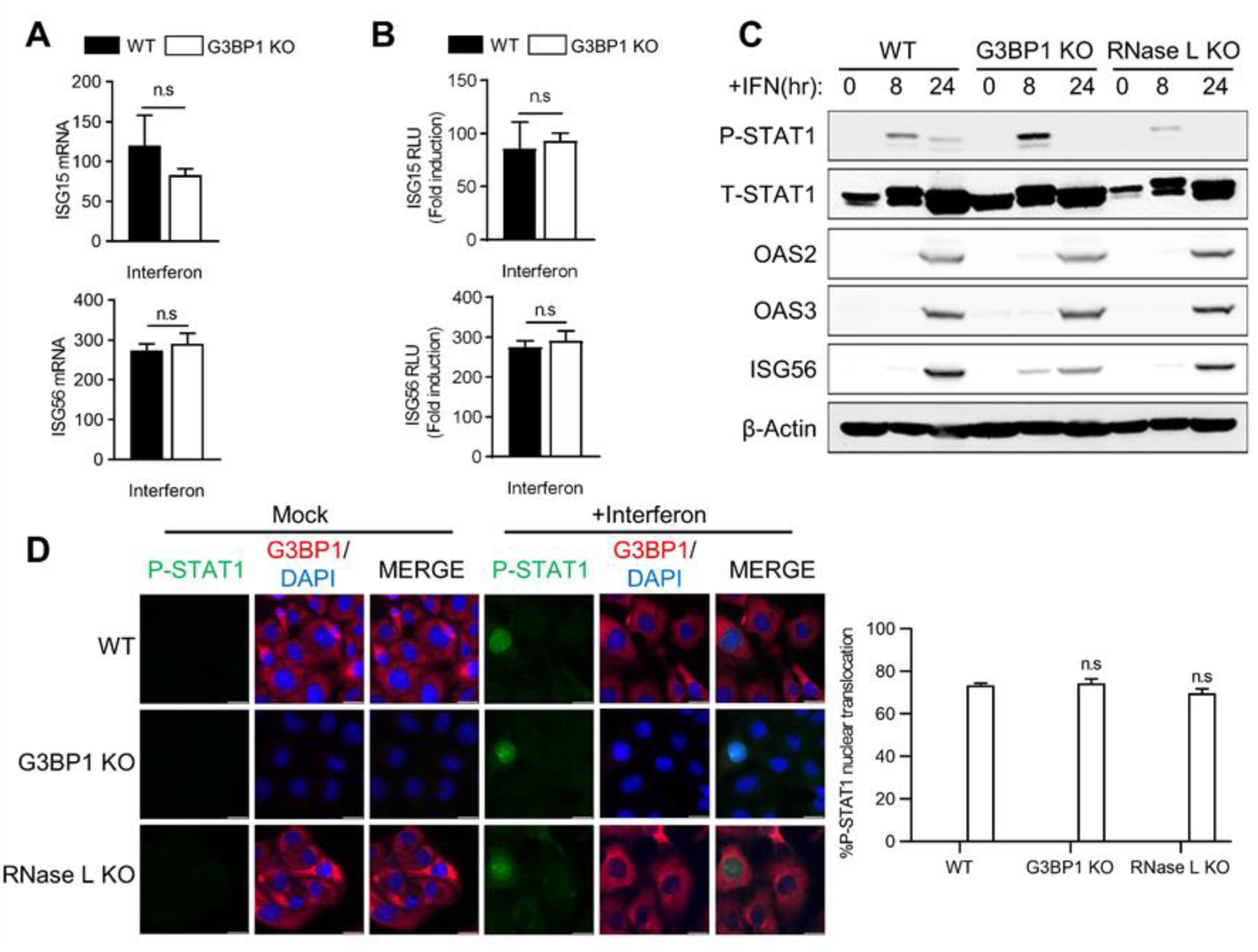
Effect of avSG formation on IFN signaling. HT1080 WT and G3BP1 KO cells were (A) treated with IFN-β (1000 U/ml) for 24h and mRNA levels of ISG15 and ISG56 were measured and normalized to GAPDH by qRT-PCR, (B) transfected with ISG15-luc or ISG56-luc reporter constructs along with β-galactosidase plasmids and 24h later treated with IFN-β (1000 U/ml) and luciferase activity were measured and normalized to β-galactosidase levels. (C) WT, G3BP1 KO and RNase L KO cells were treated with IFN-β (1000 U/ml) for indicated times and cell lysates were analyzed for phosphorylation of STAT1 and induction of OAS2, OAS3 and ISG56 in immunoblots. β-actin was used to normalize loading. (D) WT, G3BP1 KO and RNase L KO cells were treated with IFN-β (1000 U/ml) for 16h and nuclear translocation of p-STAT1 was determined by immunofluorescence and nucleus was stained with DAPI, (right) quantification of p-STAT1 nuclear translocation from five random fields. Data represent mean ± S.E. for three independent experiments. n.s: not significant, WT: Wild-Type.

### Induction of proinflammatory cytokines by RNase L is independent of antiviral stress granule assembly

Activation of RNase L or treatment with RNase L-cleaved RNAs induces inflammatory signaling pathways and proinflammatory cytokines (47, 54). We have demonstrated that avSG is required for RNase L-mediated IFN production, however, the requirement of avSG in inducing proinflammatory cytokines during RNase L activation is not known. To determine the effect of avSG on cytokine induction during RNase L activation, we monitored CCL5 (RANTES), IL-8 or IP-10 promoter activation using luciferase reporter constructs in WT and G3BP1 KO cells by directly activating RNase L with 2-5A or introducing RNase L-cleaved small RNAs and compared to control small RNAs. The 2-5A-induction of CCL5 (RANTES), IL-8 or IP-10 promoter in G3BP1 KO cells was comparable to that of control WT cells (Fig. 6A). Consistent with the observation that RNase L-cleaved small RNAs promoted inflammasome signaling, we observed increase in promoter activation in cells treated with these RNAs compared to control small RNAs. As with 2-5A treatment, depleting G3BP1 in cells did not affect induction of CCL5 (RANTES), IL-8 or IP-10 promoter by RNase L-cleaved small RNAs in promoter-driven luciferase assays (Fig. 6B). Similar increase in mRNA levels of CCL5 (RANTES), IL-8 or IP-10 as well as CXCL1 was observed in response to 2-5A and RNase L-cleaved RNAs in control WT cells and depletion of G3BP1 did not affect mRNA levels as determined by real-time PCR analysis (Fig. 6C, D). To further analyze if Tumor Necrosis Factor alpha (TNFα)-induced cytokines are affected by SGs, we compared CCL5 (RANTES), IL-8 or IP-10 promoter induction in response to TNFα in G3BP1 KO and control WT cells. No significant difference in promoter induction of TNF-α-induced cytokines was observed in cells lacking G3BP1 (Fig. 6E). These results show that while RNase L activation and the RNA cleavage products induce proinflammatory cytokines, unlike IFN-β production, avSG induced by RNase L is not required for this effect as cells lacking key SG protein, G3BP1, induce comparable levels of these cytokines.

**Figure 6.**
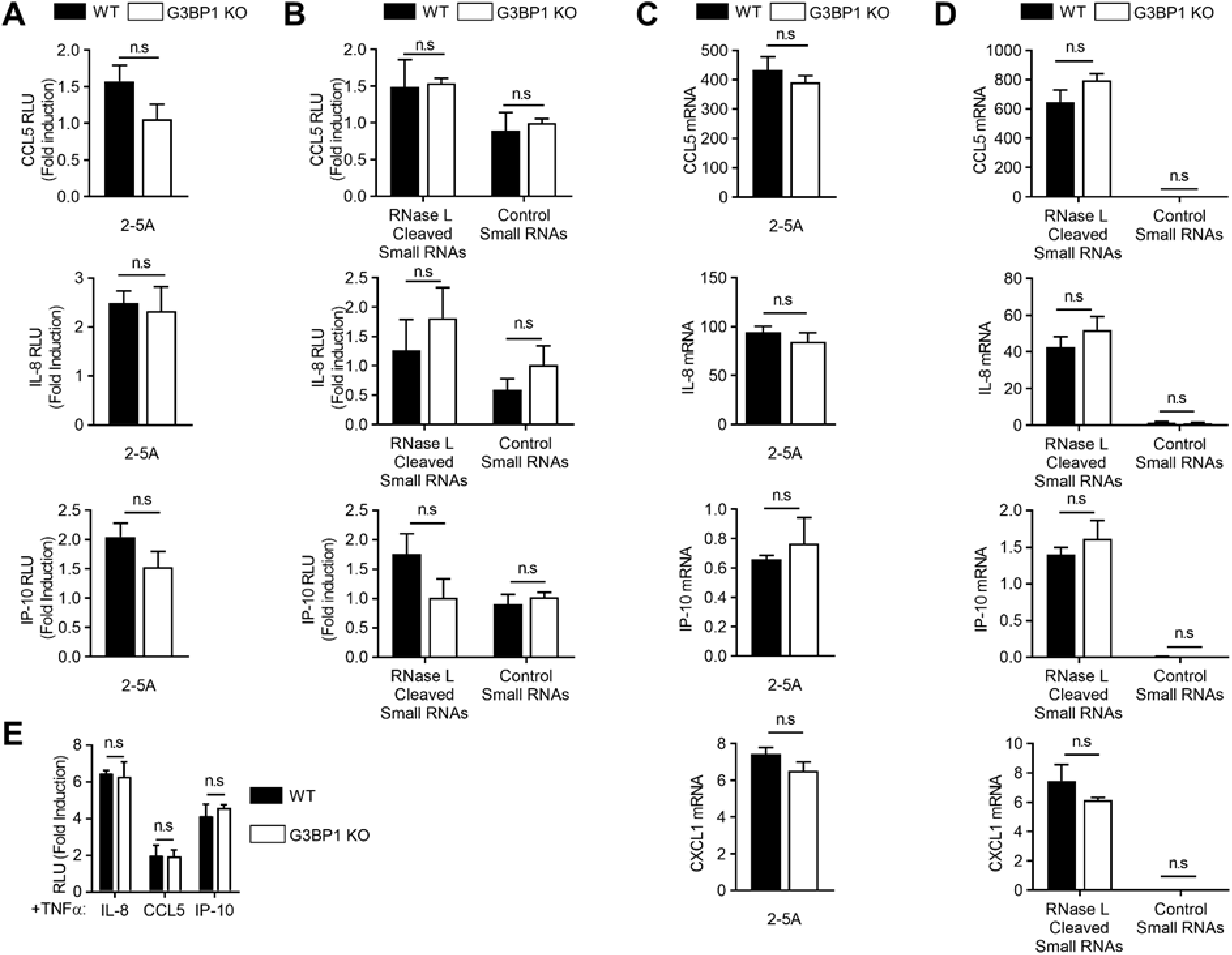
Induction of proinflammatory cytokines by RNase L is independent of avSG assembly. WT and G3BP1 KO cells were transfected with CCL5-luc, IL-8-luc or IP-10-luc and β-galactosidase plasmids and 24h later treated with (A) 2-5A (10µM), (B) RNase L-cleaved small RNAs or control small RNAs and luciferase activity normalized to β-galactosidase levels. WT and G3BP1 KO cells were transfected with (C) 2-5A (10µM), (D) RNase L-cleaved small RNAs or control small RNAs and mRNA levels of CCL5, IL-8, IP-10 and CXCL1 was measured by qRT-PCR and normalized to GAPDH mRNA levels, (E) WT and G3BP1 KO cells were transfected with CCL5-luc, IL-8-luc or IP-10-luc and β-galactosidase plasmids and 24h later treated with 100ng/ml of TNFα and luciferase activity normalized to β-galactosidase levels. Data represent mean ± S.E. for three independent experiments. n.s: not significant, WT: Wild-Type.

### Antiviral stress granule assembly restricts SeV replication

RNase L contributes to IFN-β production during Sendai virus (SeV) infection and SeV is susceptible to RNase L antiviral effects (46). We tested the hypothesis that SeV infection induces avSG formation with antiviral roles in infected cells. Virus infected cells were detected by immunostaining using anti-SeV antibodies for structural proteins 24h post infection and SG formation was monitored by appearance of G3BP1 puncta (Fig. 7A). To biochemically characterize the SG formed during SeV infection as avSG, we purified avSG core from infected cells and co-immunopreciptated antiviral proteins that interacted with G3BP1 and compared to uninfected cells. As with avSG formation in RNase L-activated cells, G3BP1 interacted with Rig-I and PKR in avSG core only during infection and OAS and RNase L localized to avSG but did not interact with G3BP1(Fig. 7B). We blocked formation of avSG to demonstrate the significance during SeV infection using cells lacking G3BP1, a protein critical for avSG assembly. To further understand the role of RNase L-induced avSG, we used RNase L KO cells and compared SeV RNA copies produced during the time course of SeV infection up to 36h. In G3BP1 KO cells increase in SeV RNA copies was observed at 24h and further increased to 2.8-fold by 36h compared to control WT cells (Fig. 7C). Consistent with previous studies, RNase L KO cells were more permissive to SeV replication and SeV RNA copies were 5-fold more at 24h and a log higher 36h post infection (Fig. 7D). Increase in viral titers in both G3BP1 and RNase L KO cells correlated with increased accumulation of SeV proteins during time course of infection on immunoblots probed with anti-SeV antibodies (Fig. 7E). The increase in viral titers correlated with decrease in IFN-β produced during SeV infection in G3BP1 KO and RNase L KO cells demonstrating the importance of both antiviral role of RNase L as well as avSG assembly in SeV replication (Fig. 7F, G).

**Figure 7.**
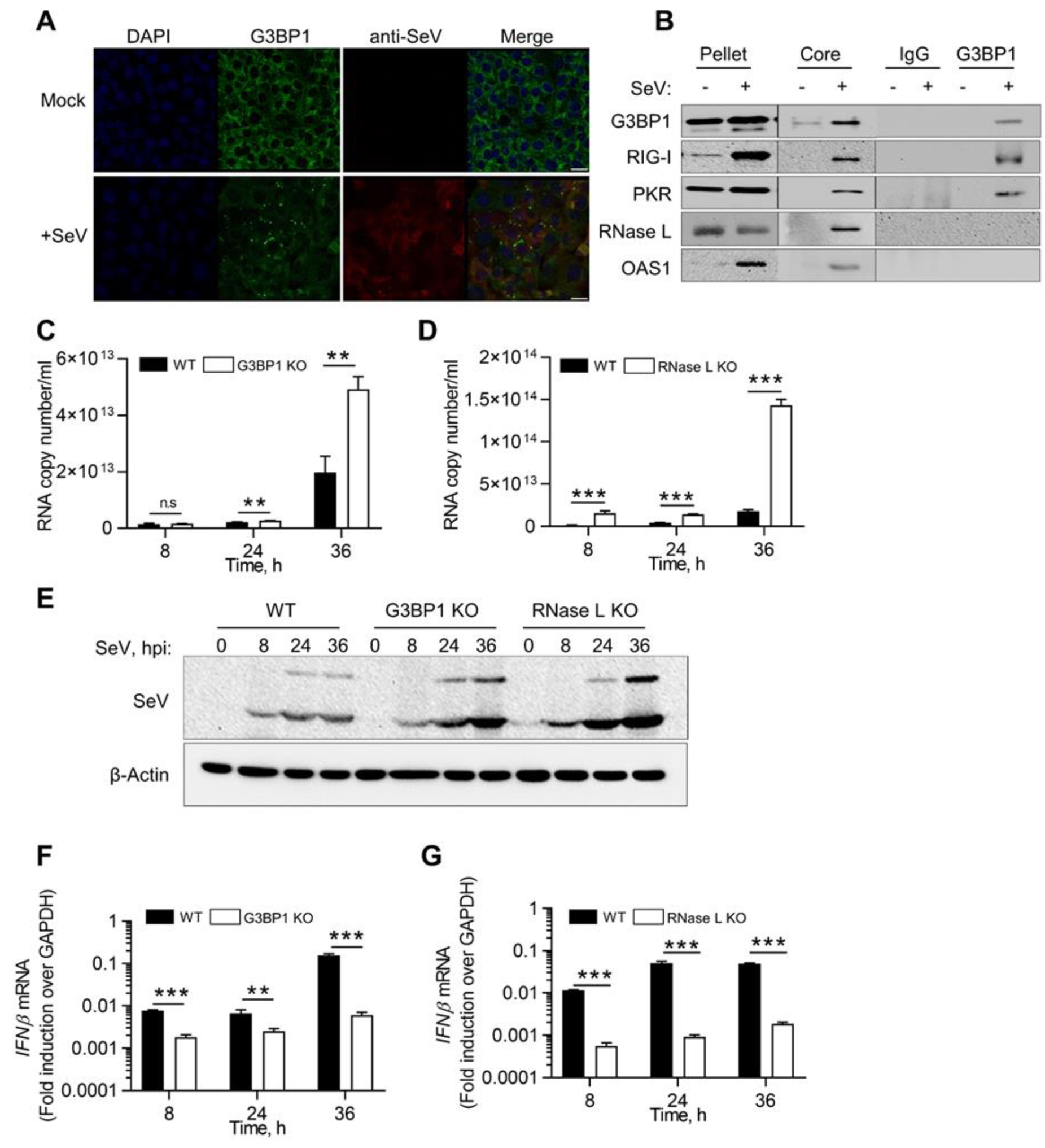
Antiviral roles of RNase L and G3BP1 during SeV infection. WT cells were infected with SeV (40 HAU/ml) for 24h and (A) cells were fixed and stained with G3BP1 and antibody against SeV, (B) avSG was purified as described in methods. The avSG core proteins were immunoprecipitated with G3BP1 antibody and immune complex analyzed for presence of PKR, Rig-I, OAS and RNase L by immunoblot analysis. Pellet and SG core fractions were probed for expression of G3BP1 and PKR, Rig-I, OAS and RNase L. Nonspecific lanes were cropped to generate the image and the boundaries are indicated. Data are representative of results from two experiments. WT, G3BP1 KO or RNase L KO cells were infected with SeV (40HAU/ml) for indicated times and (C, D) viral titers were estimated by determining copy numbers of SeV genomic RNA strands in supernatants by qRT-PCR, (E) Expression of SeV antigens were detected using anti-Sendai-virus antibody, and (F, G) IFN-β mRNA levels were measured by qRT-PCR and normalized to GAPDH mRNA levels. Data represent mean ± S.E. for three independent experiments. **p<0.001, ***p<0.0001, n.s: not significant, WT: Wild-Type.

## DISCUSSION

RNase L is a regulated endoribonuclease that is activated in virus-infected cells by a unique ligand, 2-5A (p*x*5′A(2′p5′A)*n*; *x* = 1–3; *n* ≥ 2), to produce cleavage products which are predominantly double-stranded with 5’ hydroxyl and 2’,3’-cyclic phosphate ends (55). RNase L-cleaved dsRNA activate signaling pathways by binding to diverse RNA-binding proteins to induce IFN-β, activate inflammasome, induce autophagy or promote switch from autophagy to apoptosis. Previous studies showed that RNase L cleavage products amplify IFN-β production through Rig-I and or MDA5 via MAVS (IPS-1) signaling pathway to sustain antiviral response, but how the cells coordinate RNA sensing to signaling response remains unclear (46). Our results show that RNase L activation induces antiviral stress granules (avSGs) containing key stress granule protein, G3BP1 and antiviral dsRNA binding proteins Rig-I, PKR, OAS as well as RNase L which are distinct from canonical SGs formed during oxidative stress (56). Using Crispr/Cas9 knockout cells our data suggests these dsRNAs products activate PKR and subsequent phosphorylation of eIF2α induces avSGs consistent with accumulation of dsRNA in SG with G3BP1 in response to 2-5A. Biochemical analysis of avSG using purified SG revealed interaction of G3BP1 with Rig-I and PKR, which is consistent with avSG assembled in response to virus infection and dsRNA (35, 36, 39–41). OAS and RNase L, while present in avSG core, do not physically interact with G3BP1. Finally, we demonstrate the unique requirement of avSG assembly during RNase L activation for IRF3-mediated IFN-β induction but not IFN signaling or induction of proinflammatory cytokines. Consequently, cells lacking avSG (G3BP1 KO) or RNase L signaling (RNase L KO) produced significantly less IFN during SeV infection and much higher viral titers due to compromised antiviral response. We propose that during viral infection, RNase L contributes cleaved dsRNAs to induce avSG that anchor antiviral dsRNA-binding proteins to provide a platform for efficient interaction of RNA ligands with pattern recognition receptors like Rig-I to enhance IFN-β production and antiviral response.

In our study, we have transfected cells with 2-5A, a specific ligand to directly activate RNase L and monitored formation of unique SG described as avSG based on the recruitment of dsRNA-binding antiviral proteins like Rig-I, PKR, OAS and RNase L. RNase L enzyme activity was required for avSG formation as RNase L KO cells reconstituted with functional enzyme restored avSG formation while mutant RNase L that lacked nuclease activity did not. Similar to other reports, oxidative stress by H_2_O_2_ treatment induced canonical SG formation that did not recruit antiviral proteins (56, 57). RNase L cleaves single-stranded viral and cellular RNAs after UU or UA residues leaving 5’-hydroxyl and 2’,3’-cyclic phosphate termini on dsRNAs which are required for IFN induction (46). PKR was activated by RNase L-cleaved RNAs by phosphorylating eIF2α which in turn induced avSG formation. PKR KO cells lacked phospho-eIF2α in response to RNase L-cleaved RNAs which correlated with lack of avSG formation, while cells lacking Rig-I had no effect. These results suggest that PKR is required for nucleation of avSG by RNase L. Introducing RNA cleavage products into RNase L KO cells restored avSG formation similar to control WT cells providing further evidence that RNase L-cleaved RNAs are inducers of avSG by activating PKR. Removal of the 2’,3’-cyclic phosphate termini which was required for IFN induction, decreased avSG formation demonstrating correlation of avSG formation and IFN inducing abilities. Also, dsRNA accumulated and co-localized with G3BP1 in SGs in cells treated with 2-5A. These results are consistent with avSG formed in response to IAVΔNS1, NDV, EMCV, SINV, adenovirus and Hepatitis C virus infection (26, 39, 58, 59). In other studies, formation of avSG was also observed following transfection with synthetic dsRNA, polyI:C which has broader effect by binding PKR, Rig-I or OAS isoforms (60). Binding OAS results in 2-5A production from cellular ATP that is the ligand for RNase L (10).

A recent report showed formation of unique RNase L-dependent bodies (RLB) distinct from SG in cells treated with polyI:C (61). The RLB they identify is distinct from avSGs we observe in that RLBs were formed with polyI:C treatment in cells lacking G3BP1 and was independent of SG, and did not require PKR or phosphorylation of eIF2α which were essential in our study for avSG formation. Also, the study did not explore if antiviral proteins localized with RLBs they observed. These differences may be attributed to the use of polyI:C that can bind and activate other dsRNA-binding proteins as described above. Furthermore, response to polyI:C varies in cell-type dependent manner, levels of OAS isoforms as well as abundance of polyI:C-binding proteins in cells (62). Recent reports show the role of RNase L in widespread mRNA degradation and translation repression of select basal mRNAs while antiviral mRNAs escaped decay and robustly translated (63, 64). These results suggest that RNA signaling and decay pathways activated by RNase L are complex and the dyamics may vary based on specific activation of RNase L by 2-5A compared to indirect activation by polyI:C as well as cell type differences and abundance of dsRNA-binding proteins including OAS isoforms.

We characterized the biochemical nature of avSG formed during RNase L activation using 2-5A, RNase L-cleaved RNAs and SeV infection by adapting a recently published SG purification method and determined interaction among proteins recruited to avSG. Our studies avoided overexpression of G3BP1 which forms SG independent of stimulus by testing interaction with endogenous G3BP1 (51). Interestingly, only PKR and Rig-I interacted with G3BP1 while OAS and RNase L localized but did not interact. Recent studies have shown that mature stress granule cores recruit a shell that generates a liquid-liquid phase separation from the cytosol and forms a scaffold dominated by weak RNA-protein interactions (65). Further detailed analysis will be required to determine if OAS and RNase L are present in the shell that is dynamic while PKR and Rig-I interact with G3BP1 in the inner stable core. Several other RNA helicases like DHX36, DDX3, DDX6 and antiviral proteins like ADAR1, ZAP, cGAS and Trim25 localize in avSG suggesting crosstalk between stress, RNA signaling and antiviral pathways. Future studies will address the recruitment of these additional proteins and RNA ligands in avSG and their relevance during broad range of viral infections.

Cellular and viral RNA cleavage products generated by RNase L signal to IFN-β gene through Rig-I/MDA5/MAVS (IPS-1) and IRF3 signaling pathway and here we showed involvement in inducing avSG. We demonstrate requirement of G3BP1, and thereby avSG, in IRF3 activation and IFN production using G3BP1 KO cells. RNA cleavage products are primarily responsible for avSG formation and IFN-β induction as introduction into RNase L KO cells induced IFN-β while control RNAs had no effect. In response to viral infection, activated Rig-I interacts with MAVS (IPS-1) and is redistributed on mitochondria (17, 66). Accordingly, MAVS (IPS-1) did not localize in avSG in our study consistent with similar lack of co-localization of Rig-I-containing avSG with MAVS aggregates following IAVΔNS1 infection (35). Overexpression of MAVS (IPS-1) activated downstream signaling to activate IRF3 and induced IFN-β independent of G3BP1 (Fig. 4F) and did not induce avSG indicating avSG functions upstream of MAVS-signaling and IRF3 activation. Other stress-induced pathways, like oxidative stress, do not induce avSG formation or signaling events as we have shown leading to IFN production further demonstrating the distinct nature of avSG and signaling pathways activated. AvSG assembly is not required for IFN-signaling as IFN treatment induced ISG transcription in cells lacking G3BP1 like control WT cells. No difference in nuclear translocation of phospho-STAT1 that is required for type I IFN signaling was observed in G3BP1 KO cells compared to control WT cells consistent with the role of avSG as a scaffold to recruit RNA sensors and PAMPs for signaling. RNase L also induces proinflammatory cytokines and unexpectedly, cells lacking G3BP1 induced similar levels of proinflammatory cytokines in response to 2-5A and RNase L-cleaved RNAs. Induction of cytokines by TNFα was also unaffected by lack of G3BP1. Consistent with our data, in prior studies, RNase L-cleaved RNAs stimulated NLRP3 complex formation and inflammasome activation to produce IL-1β by binding RNA helicase DHX33 and MAVS (IPS-1). Inflammasome activation was dependent on 2’,3’-cyclic phosphate termini on these RNAs and independent of both Rig-I and MDA5, but required MAVS (IPS-1) (47). Taken together, these results show a unique requirement of avSG for IRF3-mediated IFN production distinct from proinflammatory cytokines. Both studies demonstrate bifurcation of RNA-signaling pathways for proinflammatory cytokines from IFN production and appear to be independent of Rig-like receptors but dependent on MAVS (IPS-1). In other studies, overexpression of GFP-G3BP1 in HeLa and U2OS cells induced SGs localizing innate immune proteins and regulated transcription through NF-kB and JNK along with expression of cytokines (41). It is not clear if RNA ligand-induced avSG formation differ from overexpression of G3BP1 and if specific recruitment of PRRs result in specific activation of interferon vs other cytokines. Further detailed analysis of the biochemical features of the RNA ligands and the receptors will clarify how these pathways are specifically activated.

RNase L contributes to IFN-β production in vivo during SeV infection (46). Similar to RNase L KO cells, we observed reduced IFN-β production in cells lacking G3BP1 during SeV infection. And, SeV infection induced SG formation which we characterized as avSG following purification and recruitment of PKR, Rig-I, OAS and RNase L (Fig. 7A, B). Reduced levels of IFN-β production facilitated higher replication of SeV in both RNase L KO and G3BP1 KO cells with loss of antiviral effect. A recent study suggests that G3BP1 inhibits SeV and VSV replication by suppressing RNF125-mediated ubiquitination of Rig-I resulting in increased Rig-I expression and IFN production (53). These observations suggest that G3BP1 may regulate host response to viral infections at multiple levels by regulating activity of PRRs like Rig-I as well as nucleating avSG formation. Prior studies showed that SeV infection produces highly structured dsRNA copy-back intermediates (defective viral genomes, DVG) with enhanced immunostimulatory activity (67). DVGs bind Rig-I and trigger expression of type I IFN and proinflammatory cytokines in infected cells (68). Another report identified unusual RNA species produced by various SeV strains with IFN-inducing abilities that correlated with SG-like structures (69). While not explored in this study, we speculate that highly structured RNA motifs present in DVGs are released by RNase L activity similar to RNA PAMPs produced from the 3’ region of HCV genome during HCV infection to sustain IFN production (70).

RNase L has antiviral effects against broad range of RNA and DNA viruses. We demonstrate a role for RNase L-cleaved RNAs in inducing stress granules to serve as an antiviral signaling hub by coordinating interaction of RNA ligands with pattern recognition receptors (PRRs) to amplify IFN production and effectively mount antiviral response. It is likely that RNase L-cleaved RNAs are eventually turned over in P-bodies harboring mRNA decay machinery to prevent sustained activation. Further studies will evaluate the role of RNase L-induced avSG in pathogenesis of viruses susceptible to RNase L antiviral effect and the broader impact on virus infection. Also, viruses antagonize host response and SG assembly to promote replication in host cells. Critical balance of host stress response pathways and viral manipulation of these pathways eventually dictates the outcome of viral infections.

## MATERIALS AND METHODS

### Chemicals, reagents and antibodies

Chemicals, unless indicated otherwise, were from Sigma Aldrich (St. Louis, MO, USA). Antibodies against G3BP1 (SC-81940), OAS1 (SC-98424), OAS2 (SC-374238), RNase L (SC-22870), PKR (SC-707) and RIG-I (SC-48931) were from Santa Cruz Biotechnology; G3BP1 (A302-033A) used for immunoprecipitation was from Bethyl laboratories; Sendai virus (MBL-PD029) was from MBL; FLAG (14793), phospho-eIF2α (3398), eIF2α (5324), Histone H3 (9715), Lamin A/C (4777), phospho-IRF3 (4947), IRF3 (4302), phospho-STAT1 (9167), STAT1 (9172), ISG56 (14769) and β-actin (3700) were from Cell Signaling Technology; Recombinant human TNF-α (PHC3011) and antibody against OAS3 (PA5-31090) was from Thermo Fisher Scientific; RIG-I (ALX-804-849-C100) and MAVS (ALX-210-929-C100) were from Enzo Life Sciences; phospho-PKR (AB81303) and total-PKR (1511-1) were from Abcam; monoclonal antibody to human RNase L was kindly provided by Robert Silverman (Cleveland Clinic); dsRNA (J2) was from English & Scientific Consulting. Anti-mouse IgG and anti-rabbit IgG HRP linked secondary antibodies were from Cell Signaling Technology and ECL reagents were from Boston Bioproducts and GE Healthcare. Interferon β was from Biogen Idec. Hydrogen peroxide (H325-100) and puromycin (BP2956100) was purchased from Fisher scientific. 2–5A was prepared enzymatically from ATP and recombinant 2–5A synthetase (a generous gift from Rune Hartmann, University of Aarhus, Aarhus, Denmark) as described previously (71).

### Cell culture and transfections

The human fibrosarcoma cell line, HT1080 (a gift from Ganes Sen, Cleveland Clinic, Cleveland, OH, USA) were cultured in Dulbecco’s modified minimal essential medium with 10% fetal bovine serum,100 µg/mL penicillin/streptomycin, 2 mM L-glutamine, and non-essential amino acids. Cells were maintained in 95% air, 5% CO2 at 37°C. Transfection of 2-5A (10µM) was performed using lipofectamine 2000 (Invitrogen, Carlsbad, CA, USA) according to manufacturer’s protocol. RNase L cleaved small RNAs and control small RNAs were prepared as previously (46, 72) and transfected (2µg/ml) using Polyjet reagent (SignaGen Laboratories) according to manufacturer’s protocol. H_2_O_2_ (1µM) was added to cell culture media for 3 hours to induce oxidative stress.

### Generation of cells with *PKR*, *Rig-I*, *RNase L* and *G3BP1* knockout using CRISPR/Cas9 system

Knockout cells were generated using CRISPR/Cas 9 system (72, 73). Small guide RNAs (sgRNA) (Table 1, supplementary data) were designed using (8) https://portals.broadinstitute.org/gpp/public/analysis-tools/sgrna-design. The guide RNA sequences were synthesized as DNA oligonucleotides and annealed, phosphorylated and ligated into the vector pSpCas9(BB)-2A-Puro (PX459 Addgene plasmid #62988) V2.0 (a gift from Feng Zhang) that was prepared by digestion with BsmBI. HT1080 cells (3×10^5^ cells/well of a 6-well plate) were transfected with 2µg resulting plasmids and selected in (1µg/ml) puromycin. Clones were obtained by limiting dilution and gene knock-out colonies were validated by immunoblot and sequencing (Fig S1).

**Table 1:**
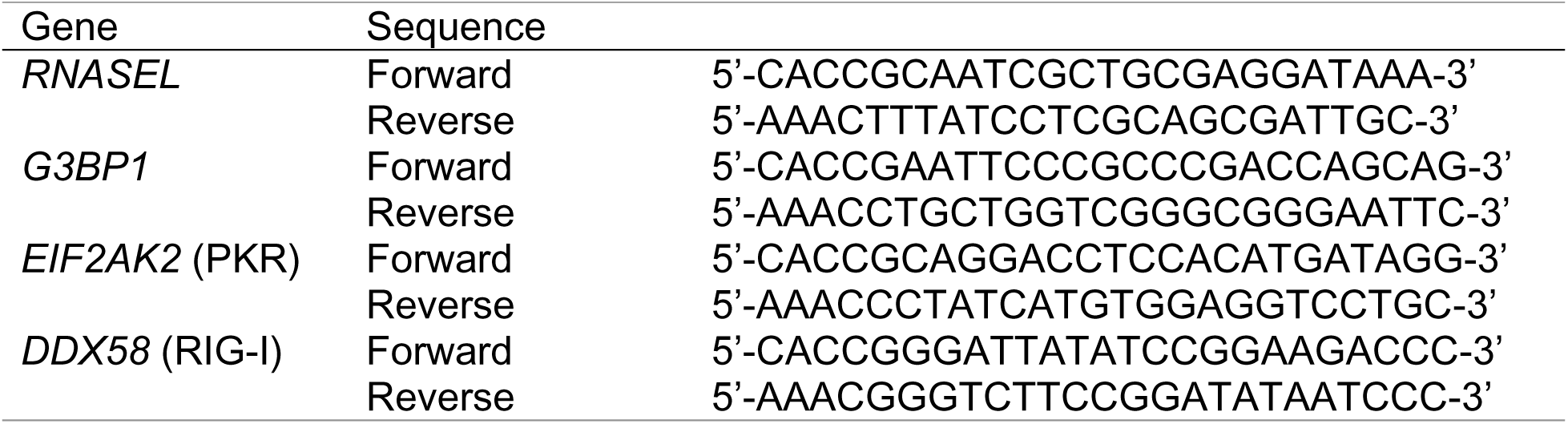
sgRNA sequence for *RNase L, G3BP1, PKR* and *Rig-I* knockout using CRISPR/Cas9 system.

### Plasmids

Plasmids Flag-RNase L, Flag-RNase L R667A (Robert Silverman, Cleveland Clinic), GFP-IRF3 (Travis Taylor, University of Toledo), HA-IPS-1(MAVS) (Invivogen), IFN-β-luc (Michael Gale, University of Washington), ISG56-luc (Ganes Sen, Cleveland Clinic), ISG15-luc (Bret Hassel, University of Maryland), IP10-luc, IL-8-luc (George Stark, Cleveland Clinic), CCL5-luc, IRF3-Gal/UAS-luc (Katherine Fitzgerald, University of Massachusetts) were transfected using Polyjet reagent as per manufacturer’s instructions.

### Western blot analysis

The cells were lysed in NP-40 lysis buffer containing 0.5% NP-40, 90 mM KCl, 5 mM magnesium acetate, 20 mM Tris, pH 7.5, 5 mM β-mercaptoethanol, 0.1 M phenylmethylsulfonyl fluoride (PMSF), 0.2 mM sodium orthovanadate, 50 mM NaF, 10 mM glycerophosphate, protease inhibitor (Roche Diagnostics). The lysates were clarified by centrifugation at 10,000×g (4°C for 20 min). Protein (15–100 μg per lane) was separated in polyacrylamide gels containing SDS and transferred to Nitrocellulose membrane (Biorad) and probed with different primary antibodies according to the manufacturer’s protocols. Membranes were incubated with goat anti-mouse or goat anti-rabbit antibody tagged with horseradish peroxidase (Cell Signaling) and immunoreactive bands were detected by enhanced chemiluminesence (GE Healthcare and Boston Bioproducts). Images were processed using Adobe Photoshop CS4 (Adobe, San Jose, CA, USA). In some instances, nonspecific lanes were cropped to generate the images and the boundaries are indicated in representative figures.

### Immunofluorescence analysis

Cells were cultured on glass coverslips and after treatment, cells were fixed with 4% paraformaldehyde (Boston Bioproducts) for 15 minutes and permeabilized with 0.1% Triton X-100 in PBS for 15 minutes. Cells were then blocked with 3% BSA for 1 hour at room temperature and incubated overnight at 4°C with indicated antibodies. Alexa488- or Alexa647-conjugated anti-immunoglobulin antibody (Molecular Probes) were used as secondary antibodies. Cell nuclei were stained with Vectashield with DAPI to stain the nucleus (Vector Labs). Fluorescence and confocal microscopy assessments were performed with Leica CS SP5 multi-photon laser scanning confocal microscope (Leica Microsystems). All subsequent analysis and processing of images were performed using the LAS AF software (Leica Microsystems). Cells containing avSG (n>5) which are above 0.6µm in diameter were considered for analysis. The percentage of avSG containing cells were calculated in at least five random fields from a minimum of 100 cells per treatment. Colocalization of proteins in stress granules were assessed by line scan analysis using Image J as described (74). A line was drawn across the stress granules and the intensity were measured using plot profile. The arbitrary intensity was plotted according to arbitrary distance for each channel.

### RNA isolation, rRNA cleavage assay and Quantitative real-time PCR

Total RNA was isolated from cells using Trizol reagent (Invitrogen), as per manufacture instructions and resolved on RNA chips using Bioanalyzer 2100 (Agilent Technologies) as described previously (71). RNase L cleaved small RNAs, CIP treated RNase L-cleaved small RNAs and control small RNAs were purified as described earlier (46, 71). Reverse transcription and cDNA synthesis was performed using random decamers and a RETROscript cDNA synthesis kit (Life Technologies; Thermo Fisher Scientific). Gene expression was determined by quantitative reverse transcription polymerase chain reaction (qRT-PCR) using SYBR Green PCR Master Mix (Bio-Rad Laboratories Inc., Hercules, CA, USA) using the gene-specific primers (Table 2, supplementary data) and normalized to GAPDH expression.

**Table 2:**
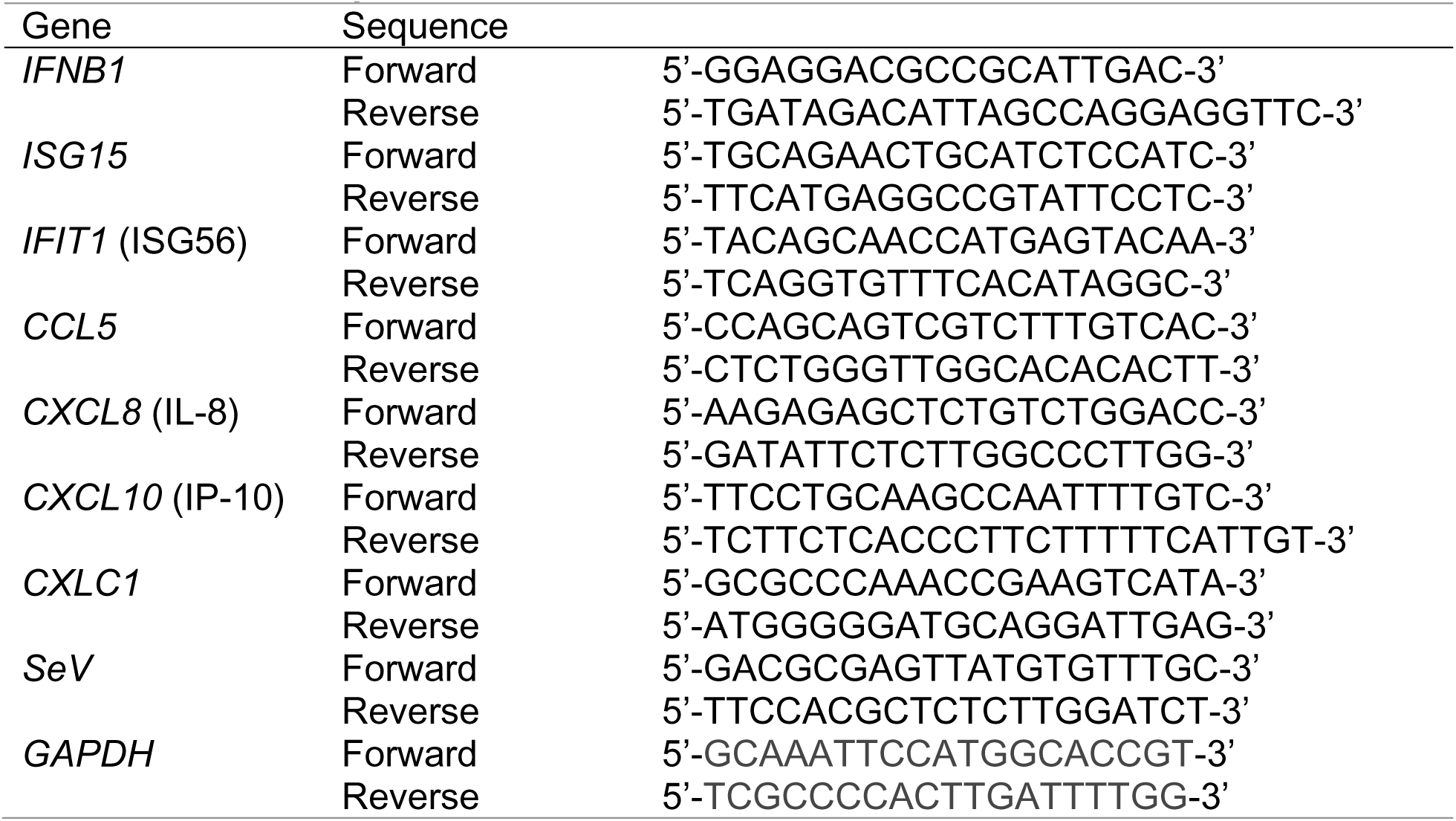
Primer sequences for real time RT-PCR.

### Luciferase assay

Cells (1×10^5^) were seeded in 12-well plate and transfected with indicated plasmids along with pCH110 β-galactosidase expressing plasmid to normalize transfection efficiency. Cells were harvested at indicated time points in luciferase lysis buffer and luciferase activity was determined using luciferase reagents (Goldbio,USA) and normalized to β-galactosidase levels (75).

### Stress granules isolation and immunoprecipitation

Stress granules were isolated as described before (51). Briefly, cells were grown on six 10cm dishes and after stress, cells were pelleted at 1500×g for 3 min. Upon removal of media, pellets were immediately flash-frozen in liquid N2 and stored at –80°C until isolation of the stress granule core was performed. Cell pellet was thawed on ice for 5 min, resuspended in 1ml of SG lysis buffer (50 mM Tris-HCl (pH 7.4), 100 mM potassium acetate, 2 mM magnesium acetate, 0.5 mM dithiothreitol, 50 µg/ml heparin, 0.5% NP-40, EDTA-free protease inhibitor, 1 U/µl of RNasin plus RNase inhibitor (Promega) and passed through a 25-gauge 5/8 needle attached to 1ml syringe 10 times. After lysis, lysates were spun for 5 mins at 1000×g at 4°C to remove cell debris. Supernatant was spun at 18,000×g for 20 mins at 4°C to pellet SG core. The resulting supernatant was discarded, and pellet was resuspended in 1ml of SG lysis buffer and spun at 18,000×g for 20 mins at 4°C. The resulting pellet was resuspended in 300µl of SG lysis buffer and spun at 850×g for 2 mins at 4°C. The supernatant which represents the SG core enriched fraction was transferred to new tube. Equal amounts of SG core was subject to immunoprecipitation using anti-G3BP1 antibody (1ug) and isotype specific control antibody and protein A-sepharose beads (Sigma-Aldrich). Samples were incubated at 4⁰C overnight on a rotator and immune complexes recovered by centrifugation and five washes in buffer. Samples were boiled in SDS-sample buffer and analyzed by protein gel electrophoresis and immunoblotting using indicated antibodies.

### Viral growth kinetics

5×10^5^ cells were plated in a 6-well plate and next day, cells were infected with Sendai virus (Cantell strain) at 40HAU/ml in media without serum. After 1 hour, media was replaced with complete media and cells were harvested at indicated time points. Expression of viral antigen was determined on western blots using anti-Sendai virus antibody. Total RNA was isolated from infected cells using TRIzol reagent (Invitrogen) or QIAmp viral RNA kit (Qiagen) and qRT-PCR was performed to quantify viral RNA copy number as described previously (71).

### Statistical analysis

All values are presented as mean ± SEM from at least three independent experiments or are representative of three independent experiments performed in triplicate and shown as mean ± SD. Student’s t-tests were used for determining statistical significance between groups using Prism8 (GraphPad) software and p<0.05 was considered significant.

## SUPPLEMENTARY MATERIALS

**Fig. S1.**
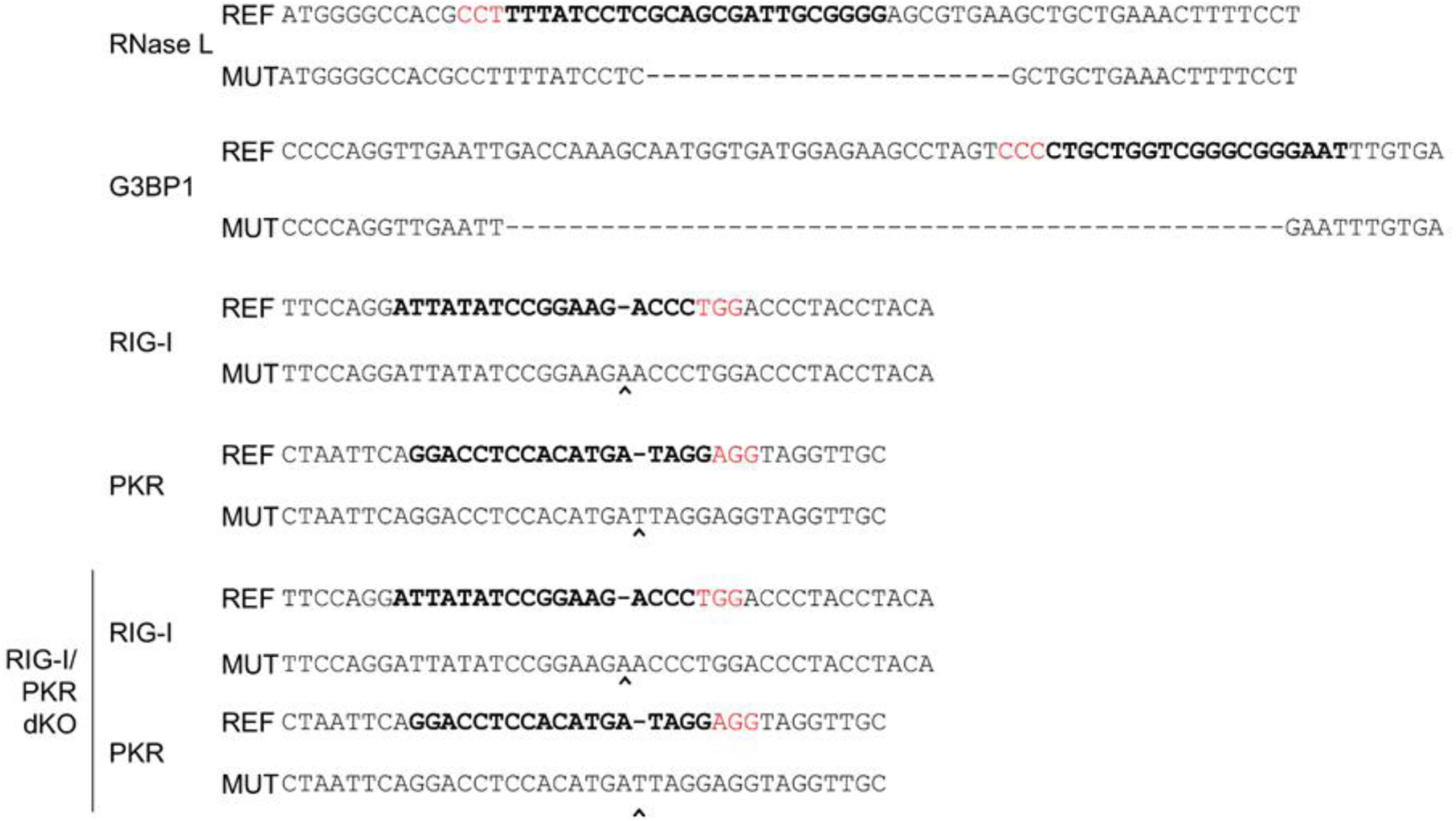
Schematic presentation of the coding regions of RNase L, G3BP1, RIG-I, PKR and RIG-I/PKR dKO that are targeted by CRISPR-Cas9. The reference sequences and the mutated sequence for each gene is shown, as confirmed by sequencing. The bold letters denote the protospacer (sgRNA binding site), the red letters indicate the PAM (protospacer adjacent motif).

## ACKNOWLEDGEMENTS

We thank Robert Silverman, Ganes Sen and George Stark (Cleveland Clinic) for providing cells, plasmids and antibodies used in this study. This work was supported by National Institutes of Health (NIH) Grants AI119980-01A1 (KM), internal grants (KM) and startup funds from University of Toledo (KM). We thank Scott Leisner (University of Toledo), Saurabh Chattopadhyay (University of Toledo) and Travis Taylor (University of Toledo) for valuable discussions through the course of this work.

